# igv.js: an embeddable JavaScript implementation of the Integrative Genomics Viewer (IGV)

**DOI:** 10.1101/2020.05.03.075499

**Authors:** James T. Robinson, Helga Thorvaldsdóttir, Douglass Turner, Jill P. Mesirov

## Abstract

igv.js is an embeddable JavaScript implementation of the Integrative Genomics Viewer (IGV). It can be easily dropped into any web page with a single line of code and has no external dependencies. The viewer runs completely in the web browser, with no backend server and no data pre-processing required.

## Introduction

The rapid pace of genomic dataset generation has led to a proliferation of public and private web portals and cloud resources for sharing, analyzing, and visualizing the data. An important component of many such resources is the ability to visually inspect the data in track-based views aligned to a reference genome. Traditionally such views have been provided in genome browsers, including large client-server systems such as the UCSC Genome Browser [1] and the Ensembl Genome Browser [2], rich client web applications such as JBrowse [3], and desktop applications such as the Integrative Genomics Viewer (IGV) [4,5]. Although it is possible to create links from web portals to visualize data in these and similar browsers, in many cases it is desirable to embed genomic visualizations directly in portal pages, enabling data visualization in place without the context switch to another web page or application. Here we describe igv.js, an embeddable JavaScript implementation of the IGV genome viewer.

## Features

Unlike stand-alone genome browsers such as JBrowse, the igv.js viewer has been designed explicitly to be embedded in and controlled by enclosing web applications. It builds on previous work, importantly Dalliance [6], the first pure JavaScript component to natively support next generation sequencing (NGS) data in binary BAM files in a web browser. Embedding an igv.js visualization is as simple as creating a container “div” element in the HTML document and calling a JavaScript function to insert the genome browser. The enclosing application interacts with igv.js by means of an API, which includes functions to configure the initial view, load and remove reference genomes and tracks, navigate to specific loci, and listen for events initiated by user interaction with igv.js. Tracks are initialized with configuration objects that define the track type, data source and format, and initial visual properties such as color, height, and feature layout within the track. Multiple instances of igv.js can exist on a page, and instances can be added and removed dynamically.

The look and feel of igv.js is modeled closely after the IGV desktop application. Like the desktop IGV application, igv.js supports a wide range of genomic track types and file formats, including aligned reads, variants, coverage, signal peaks, annotations, eQTLs, GWAS, and copy number variation (see examples in Figure 1). A particular strength of IGV is manual review of genome variants, both single-nucleotide and structural variants [7]. To support this use case in igv.js we have implemented the suite of IGV read alignment display options. These include coloring and sorting reads by strand and other attributes, displaying and coloring bases that mismatch the reference, coloring alignments to flag ambiguous mappings and anomalous pair distance and orientations, and multi-locus views for side-by-side views of the multiple components of a structural variant.

**Figure 1.**
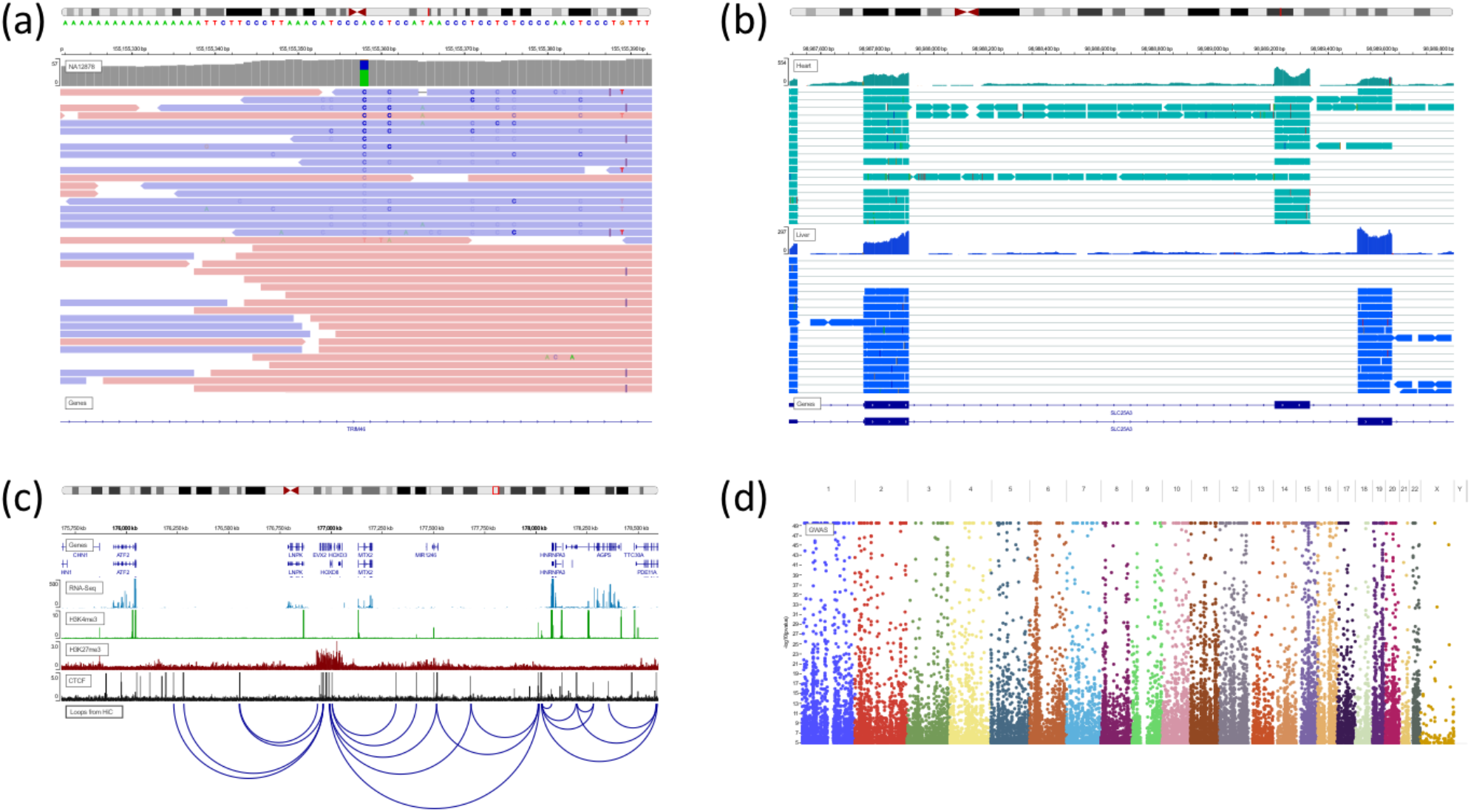
Example igv.js screenshots. (a) NGS sequence alignment pileup in the region of a putative SNP indicates strand bias (https://tinyurl.com/y9n6dyw9). (b) RNA-Seq pileups from two tissues illustrate alternative splicing (https://tinyurl.com/ybo547uf). (c) Epigenetic marks and transcription factors correlate with loop calls from HiC data (https://tinyurl.com/y75465od). (d) Manhattan plot from whole-genome association studies (https://tinyurl.com/ydhakvy4).

Supported data sources include (1) files in standard genomic formats hosted on web servers, including cloud providers such as Google Drive, Google Cloud Storage, Dropbox, and Amazon S3; (2) htsget [8] servers; (3) custom web services; (4) inline data URIs; and (5) files on the local file system. To support viewing large files, such as reference sequence FASTA, BAM alignment, and VCF variant files, indices are used to fetch only the data required to render the current view utilizing standard HTTP Range requests.

Authenticated access to restricted datasets is supported with explicit support for oAuth [9] tokens, which can be supplied by the enclosing application as a static string, a callback function, or a JavaScript promise. The support for functions and promises enables the enclosing application to update or revoke tokens as needed. Additionally, arbitrary HTTP request headers can be specified during track initialization to support other token-based authentication schemes. Finally, the URLs to track resources can be specified as a callback function or promise, allowing the enclosing application to intercept and modify the URL at runtime based on user credentials or other criteria. For example, a signed URL might be generated and supplied via a promise after user authentication. Finally, explicit support is provided for Google resources by specifying a Google client id on igv.js initialization. If specified, oAuth authentication will be initiated by igv.js as required.

A unique feature of igv.js is an API function to return a compressed URL-safe string that encodes the complete state of the browser session, including pointers to the reference genome and the loaded data, as well as details on the current view and track and browser settings. This capability can be used by applications to support bookmarks and sharable links. We take advantage of this feature in IGV-Web (https://igv.org/app), an end-user application built around the igv.js component. IGV-Web provides a user interface for specifying the reference genome, loading data into the viewer from remote and local files, and saving and loading sessions. A *Share* menu allows the user to easily retrieve a URL that represents the current session state and to share it with others. The links provided in the legend for Figure 1 were created this way. Entering them into a web browser will open the IGV-Web app and recreate the interactive sessions used to generate the images in the figure.

## Availability

The igv.js distribution consists of a single JavaScript file with no external dependencies. It can be installed from NPM at https://www.npmjs.com/package/igv. The source package is available on GitHub at https://github.com/igvteam/igv.js under a permissive open-source license (MIT) and includes numerous examples. The NPM site includes a quick-start guide, and more extensive documentation is available at https://github.com/igvteam/igv.js/wiki. The IGV-Web application is available at https://igv.org/app; the source code is available on GitHub at https://github.com/igvteam/igv-webapp.

